# Interplay between copy number, dosage compensation and expression noise in *Drosophila*

**DOI:** 10.1101/041038

**Authors:** Dong-Yeon Cho, Hangnoh Lee, Damian Wojtowicz, Steven Russell, Brian Oliver, Teresa M. Przytycka

## Abstract

Gene copy number variations are associated with many disorders characterized by high phenotypic heterogeneity. Disease penetrance differs even in genetically identical twins. Can such heterogeneity arise, in part, from increased expression variability of one dose genes? While increased variability in the context of single cell gene expression is well recognized, our computational simulations indicated that in a multicellular organism intrinsic single cell level noise should cancel out and thus the impact of gene copy reduction on organismal level expression variability must be due to something else. To systematically examine the impact of gene dose reduction on expression variability in a multi-cellular organism, we performed experimental gene expression measurements in *Drosophila* DrosDel autosomal deficiency lines. Genome-wide analysis revealed that autosomal one dose genes have higher gene expression variability relative to two dose genes. In flies, gene dose reduction is often accompanied by dosage compensation at the gene expression level. Surprisingly, expression noise was increased by compensation. This increased compensation-dependent variability was found to be a property of one dose autosomal genes but not *X*-liked genes in males despite the fact that they too are dosage compensated, suggesting that sex chromosome dosage compensation also results in noise reduction. Previous studies attributed autosomal dosage compensation to feedback loops in interaction networks. Our results suggest that these feedback loops are not optimized to deliver consistent responses to gene deletion events and thus gene deletions can lead to heterogeneous responses even in the context of an identical genetic background. Additionally, we show that expression variation associated with reduced dose of transcription factors propagate through the gene interaction network, impacting a large number of downstream genes. These properties of gene deletions could contribute to the phenotypic heterogeneity of diseases associated with haploinsufficiency.

**Author Summary:** Gene copy number variations are associated with many human disorders characterized by high phenotypic heterogeneity and understanding the effects of gene copy alterations is essential if we wish to understand factors influencing disease heterogeneity. While heterogeneous responses to reductions in gene dosage can be attributed in part to genetic background differences, we found that responses to dosage reduction vary, even in identical genetic backgrounds. Heterogeneities are not restricted to reduced dosage genes, but propagate through gene networks to affect downstream genes with normal copy number. We also found that reduction in gene dosage is associated with the introduction of expression variation or noise. Expression noise was also observed to propagate across the gene network, further contributing to the heterogeneous response to gene deletion. Through the use of computational simulation, we showed that the majority of the increased noise we observed is most likely due to extrinsic rather than intrinsic sources. Irrespective of the source of the observed noise, we propose that the presence of expression noise and its propagation through gene networks is likely to contribute to the heterogeneity of disease phenotypes observed in humans.

## Introduction

Gene expression is a multi-step process that is stochastic in nature, with apparently isogenic cells displaying cell-to-cell differences in gene expression: a phenomenon commonly referred to as expression noise [1–3]. Following pioneering work at the single cell level [4] and the first mathematical models of the process [5], we typically distinguish two major sources of stochastic gene expression variation. An intrinsic contribution resulting from stochasticity of biochemical processes, including factors such as transcriptional bursting and gene copy number; and an extrinsic component, which arises from subtle environmental differences, cell-cell communication, or randomness within cell populations, including cell size and cell-cycle phase.

Gene dose change is an important genomic structural alteration that can influence gene expression and organismal phenotype. For example, in diploid budding yeast an estimated 3% of the genome is revealed to be haploinsufficient when assayed by growth of deletion mutants in standard rich medium, and much of this effect is due to reduced expression from one copy genes [6]. Mathematical models of single cell expression noise indicate that a reduction in gene copy number increases intrinsic noise [7,8]. Intuitively, starting with the assumption that in eukaryotic organisms gene expression occurs in bursts [9,10], expression from two gene copies regulated by independent promoters leads to smaller expression noise relative to doubled expression from one promoter. It has been proposed that the observed fitness advantage that diploid yeasts have over haploids results from the reduction of expression noise by genome doubling [11].

In contrast to work with single cell organisms, the impact of gene dosage on expression variability in metazoans is less well studied. However, a full understanding of the effect of gene copy deletions is fundamental for better understanding of diseases that originate from gene copy number changes. In humans, reduction in gene dosage for many transcription factors leads to haploinsufficient developmental disorders [12]. Thus it is likely that genomic responses to alterations in gene copy number are important drivers of some human diseases and understanding these effects may have important therapeutic implications.

Based on single cell simulations, it was hypothesized that disease susceptibility could be enhanced by haploinsufficiency [8]. In a subsequent study, it was also demonstrated that haploinsufficiency for the tumor-suppressor gene Neurofibromin 1 (*NF1*) increased variation of dendrite formation in neurofibromatosis type 1 patients [13]. Yet, it was also observed that in mammals, expression variability of *X* linked genes (single copy in males and a single active copy in females as a result of *X*-inactivation) is not different from that of autosomal genes [14]. Thus, it is not clear to what extend the results from single cell experiments can be applied to understanding expression variability in multicellular organisms.

To systematically examine the impact of gene dose reduction on expression variability in a multi-cell organism, we combined experimental gene expression measurements in *Drosophila melanogaster* autosomal deficiency lines from the DrosDel collection [15,16] and computational modeling. The DrosDel collection consists of fly lines from a common genetic background that harbor engineered deletions of different chromosomal regions, leaving a few genes in each line as one copy rather than two. In the companion paper (Lee et.al) we analyze the general effect of *Df(2L)s* on the transcriptome, including expression level of one-copy genes, properties of gene dosage compensation, relationship to chromatin domains, and impact on the expression of network neighbors. Here, analyzing gene expression differences across the genome between biological replicated RNA-seq experiments, we focus on the effect of these deletions on gene expression variability. We found that one dose genes have higher gene expression variability relative to two dose genes. Since organismal level gene expression measurements average expression over millions of cells, we asked if it was possible that the observed differences in variability were caused by intrinsic gene expression noise. In contrast to previous assumptions [14], our simulations indicate that expression variation between biological replicates cannot be attributed to intrinsic gene expression noise and thus has to be a result of a factor(s) extrinsic to individual genes and impacting expression across the whole cell population.

In flies, gene dose reduction is often accompanied by dosage compensation at the gene expression level. In particular, the male-specific lethal (MSL) complex associates with, *X* linked genes in males to increase expression two-fold relative the level of each of the two *X* chromosomes in females, thus matching gene dosage between the heterogametic (*XY*) and homogametic (*XX*) sexes relative to autosomes [17–19]. In addition, reduction of autosomal gene copy number is often, but not always, accompanied by partial dosage compensation. We have previously shown [20] and subsequently confirmed in a significantly larger study (Lee et al., companion paper), that gene dosage compensation is generally locus-specific, suggesting that feedback provided through biochemical processes and regulatory circuits account for most autosomal dosage compensation. Interestingly, we found a relationship between autosomal dosage compensation and expression variation suggesting that, in the context of gene dosage change, such feedback is noisy. However, we did not observe increased expression noise for *X* linked genes in males despite the fact that these genes are normally fully compensated. Thus MSL-dependent dosage compensation suppresses regulatory noise in addition to boosts expression levels without increasing noise.

Complex diseases are best understood from the perspective of disregulated pathways rather than individual genes. We show that expression variation associated with reduced transcription factor gene copy number propagates through the interaction network and affects expression variability of downstream genes, thus contributing to the extrinsic noise downstream of their targets.

Overall, this study for the first time demonstrates a link between gene expression variation in one dose autosomal genes, dosage compensation and propagation of expression variation associated with transcription factor gene dosage through gene interaction networks. This property of gene deletion could contribute to differences in the penetrance of disease phenotypes associated with gene deletions and haploinsufficiency in humans and suggests a novel function for X chromosome dosage compensation.

## Results

### Reduction in autosomal gene dosage leads to increased expression variability

To assess the impact of reductions in gene dose on population level expression variability, we used 99 DrosDel deficiency lines that carry different deletions on the left arm of chromosome *2 (2L).* Each deletion line was outcrossed to the *w^1118^* progenitor line [15,16] and we collected both female and male hemizygous adults. We performed RNA-Seq analysis of two biological replicates from each line and expression differences between the replicates was considered as expression variability. For the analysis, we only considered genes that were expressed above our defined technical noise level in both replicates (Lee et al., companion paper). Expression variability was quantified using δ, defined as absolute differences in FPKM (fragments per kilobase per million reads) values between two replicates normalized by the average of these two values (Materials and methods). This relative difference is linearly related to the coefficient of variation and is more natural to use when only two values are compared. Interestingly, unlike the situation observed in single cell expression noise experiments, we found no statistically significant negative correlation between δ and expression level (Fig S1). Therefore, to examine quantitatively the effect of gene dose reduction on expression variability, we pooled δ values for all hemizygous genes in the entire set of deficiencies and compared the median of δ values for one dose genes with the distribution of median δ values for the same number of two dose genes that were randomly sampled from each line (Fig 1A). For both sexes, we observed that one dose genes had significantly higher expression variability than two dose genes (*P* < 0.001). Interestingly, the difference was larger in males than females, mostly due to larger expression variability of two dose genes in females. However, there are also other differences in response to gene deletion between the sexes as we describe in the accompanying paper (Lee et al., companion paper). We also performed a chromosome-wide comparison where we considered two dose genes on the same chromosome arm (*2L*), other autosomal arms (*2R, 3L, 3R*, and *4*), and genes on the *X* chromosome (Fig 1B). As expected, we confirmed that expression variability was significantly enhanced by reductions in gene dose for all pairwise comparisons (*P* < 0.01 or *P* < 0.001, Wilcoxon rank sum test). We also found that, while there are only one dose genes on the male *X* chromosome, the median of δ values is the lowest we observed. This suggests that expression variation is reduced by *X* chromosome dosage compensation. Finally, a comparison with variations in control spike-in measures (Fig 1B) confirms that the observed variability is biological rather than technical. We confirm this observation below in analyses that correlate differences in expression variability with biological factors. However, before examining this relationship further, it was necessary to identify the nature of expression variation (intrinsic versus extrinsic) caused by reductions in gene dose.

**Fig 1.**
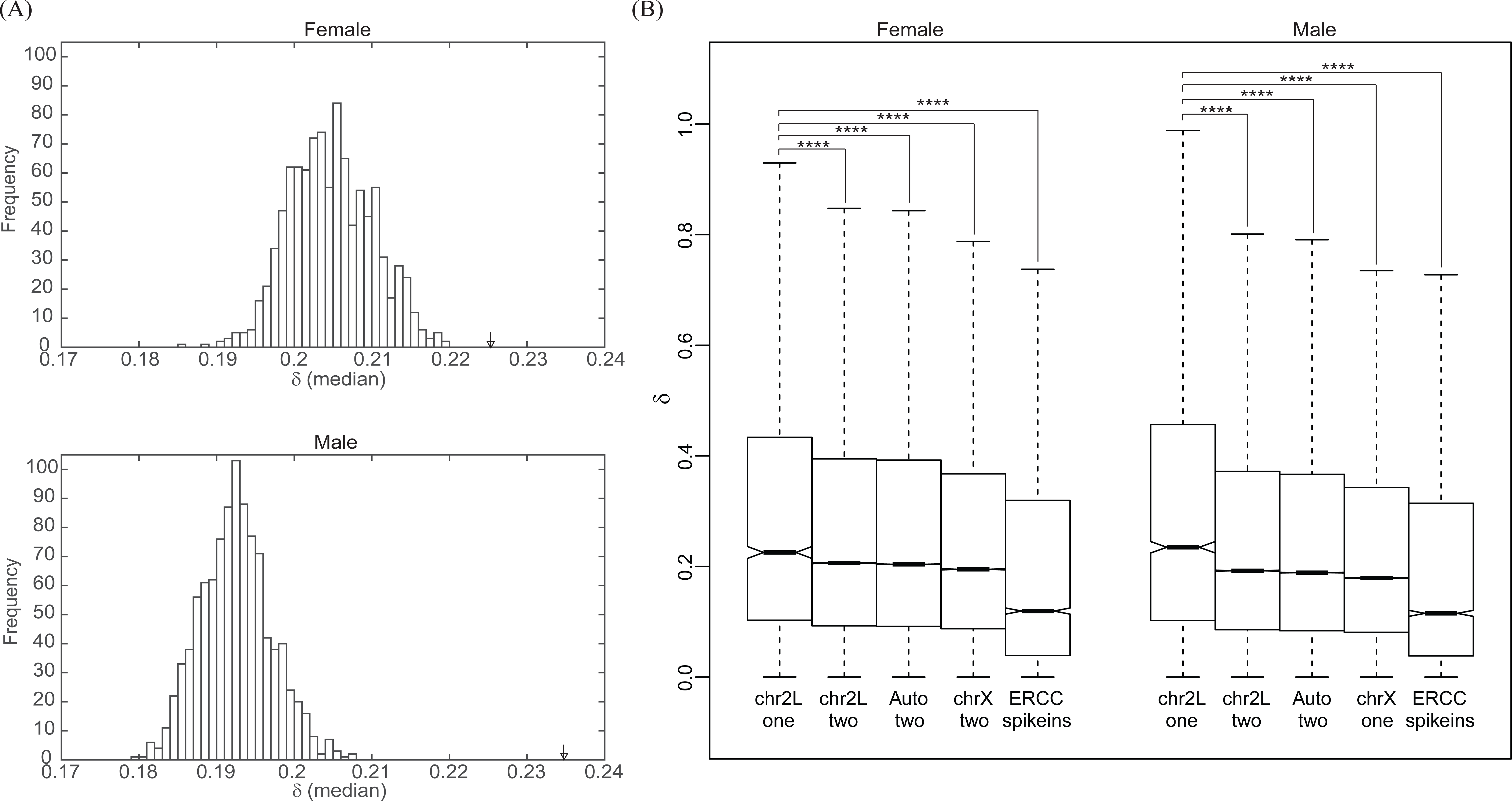
Expression variation of one dose genes. (A) The median of δ values for all one dose genes (arrow) pooled from female (above) and male (below) lines are compared with the distribution of median of δ values for the same number of two dose genes randomly sampled from each line 1000 times. (B) Boxplots show medians (bar), 95% confidence intervals (notch), 25 to 75 percentiles (box), and 1.5 × interquartile ranges (whisker) of the distribution of δ values for different copy number classes on chromosome arm *2L*, other autosomal arms, the *X* chromosome, and ERCC spike-ins for each sex (outliers are excluded). P-values were obtained from one-sided Wilcoxon rank sum tests to check whether the median of δ of one dose genes on *2L* is greater than that of two dose genes on *2L*, two dose genes on other autosomal arms, and genes on the *X*, respectively.

### Expression variation in multi-cell organisms is extrinsic in nature

In the context of single cell expression noise, theoretical models predict that if two genes are expressed at the same level but one has a larger gene copy number than the other, then the expression of the gene with the smaller number of copies is expected to be more noisy [8]. We first asked whether the increase in expression variability of one dose genes observed in the context of a multi-cell organism meets this prediction. To address this we performed stochastic simulations of gene expression in cell populations (Materials and methods). We simulated two equal cell populations to model two biological replicates. For each cell in each population, we simulated gene expression using a stochastic gene expression model (Materials and methods) and averaged the results over the cells in the population. We performed such simulations for increasing numbers of cells (Fig 2A). Our simulations indicate that if a population consists of a small number of cells, intrinsic gene expression noise can lead to very large variations in the population as measured by the δ value. However, single cell intrinsic expression noise quickly averages out for larger populations demonstrating that, at the multi-cell organism level, variability in the δ value is likely a result of extrinsic rather than intrinsic noise.

**Fig 2.**
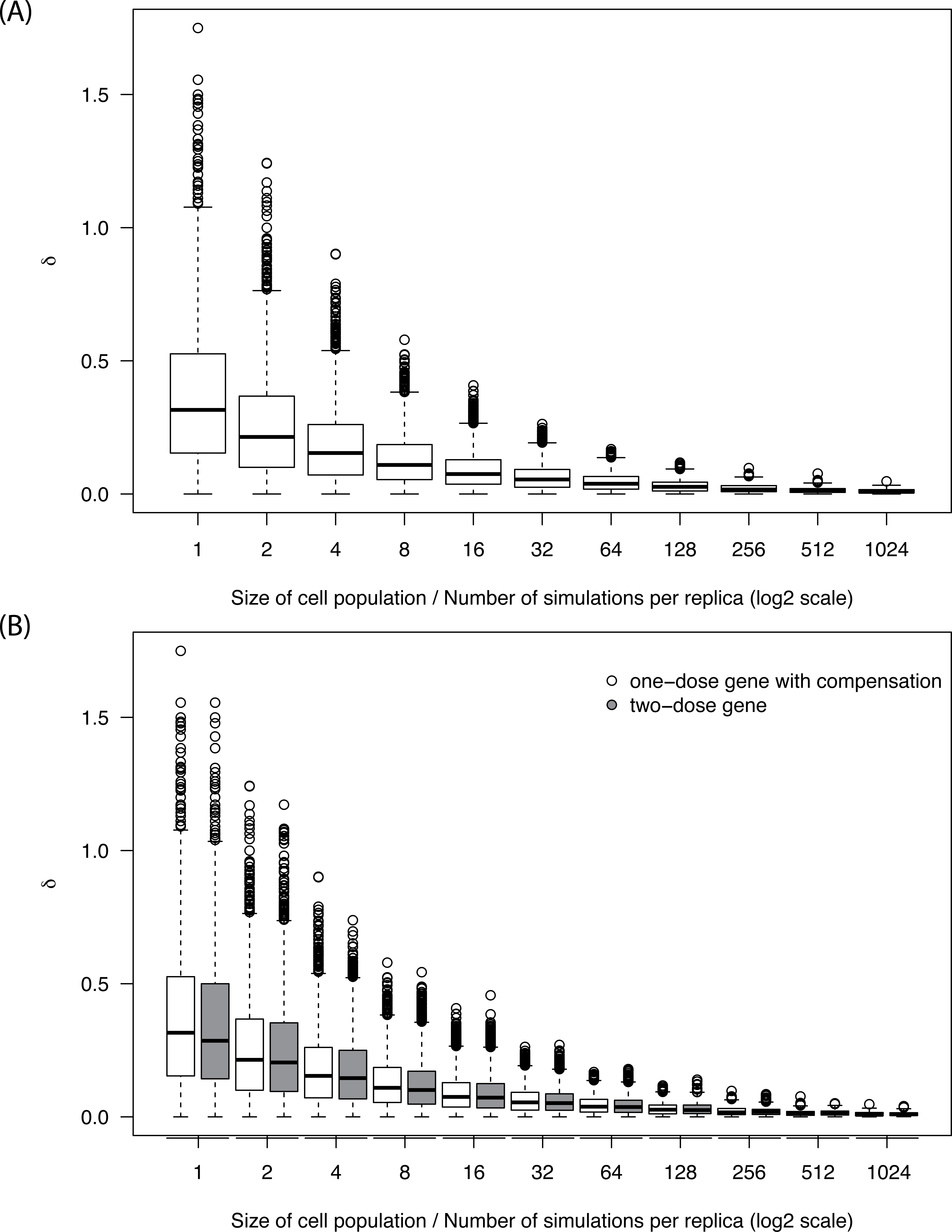
Intrinsic noise in a simulated gene expression model dissolves with increasing size of cell population. (A) Boxplots represent distribution of simulated expression variation measured by δ values for increasing size of cell population (number of simulation) per replica in a stochastic gene model. (B) Comparison of simulated variability between one dose genes (white) and two dose genes (gray) for increasing sizes of cell population. Mean gene expression is the same for both genes (full compensation). Simulations were performed using stochastic kinetic simulation of biochemical processes with the Gillespie algorithm.

We next used simulation to test whether organismal level expression variability that can arise from intrinsic noise is influenced by gene copy number reduction. To address this question, we compared the results of simulation for one and two copy genes in each cell, where mean gene expression was the same for both genes: that is, gene expression was fully compensated. As before, we performed the simulation for increasing numbers of cells and observed that the differences in expression variability due to the intrinsic component of gene copy number also disappeared in larger populations of cells (Fig 2B). These simulations demonstrated that organismal level expression variability could not be attributed to intrinsic noise, but rather must be due to extrinsic factors resulting from a more global response at the tissue-, organ-, or organism-level response.

### Relationship between expression variation and dosage compensation

Since our simulations indicated that observed expression variations of one dose genes in flies were extrinsic by nature, and since gene copy number reduction is often accompanied by some level of dosage compensation, we next asked if there was a relationship between expression variation and dosage compensation. In the case of autosomal genes, the response to reductions in gene dose is a heterogeneous gene-specific response due to gene regulatory interactions such as feedback [20]. When considering genes on the male *X* chromosome, we found that they had the smallest expression variation (Fig 1B). Similar observations have previously been made in mammals [14], although the genes examined in these cases have monoallelic expression in both males and females. This implies that MSL complex-dependent *X* chromosome dosage compensation is employed in a way that minimizes potentially harmful expression variation of *X*-linked genes in males. Indeed, using data from gene expression in female heads in the *X* chromosome deficiency lines surveyed in [21] we confirmed that the expression variation accompanying *X* chromosome deletions in females shows, similar to autosomal deletions, higher expression variability as compared to two dose genes (Fig S2). This indicates that the reduced noise is due to male-specific dosage compensation, not the particular genes found on the *X* chromosome. Thus the MSL mechanism ensures that compensation of *X*-liked genes is precise in both compensation level and expression consistency.

To test for a possible relation between autosomal dosage compensation and expression variation, we first simply grouped one dose genes according to the fold change in their expression relative to a single dosage reference (Materials and methods), then compared expression variation in each group. In the case of no compensation, we expect a two-fold expression reduction upon deletion of one copy of a gene. This is our one copy expression baseline and we refer to the variation from this baseline in any direction as a deletion response, with a positive response corresponding to dosage compensation and negative response to anti-compensation. We found that higher deletion responses (gene dosage expression compensation or larger than expected expression reduction) correspond to higher expression variations, suggesting that observed expression variation is related to the gene dosage response mediated by gene regulatory networks (Fig 3).

**Fig 3.**
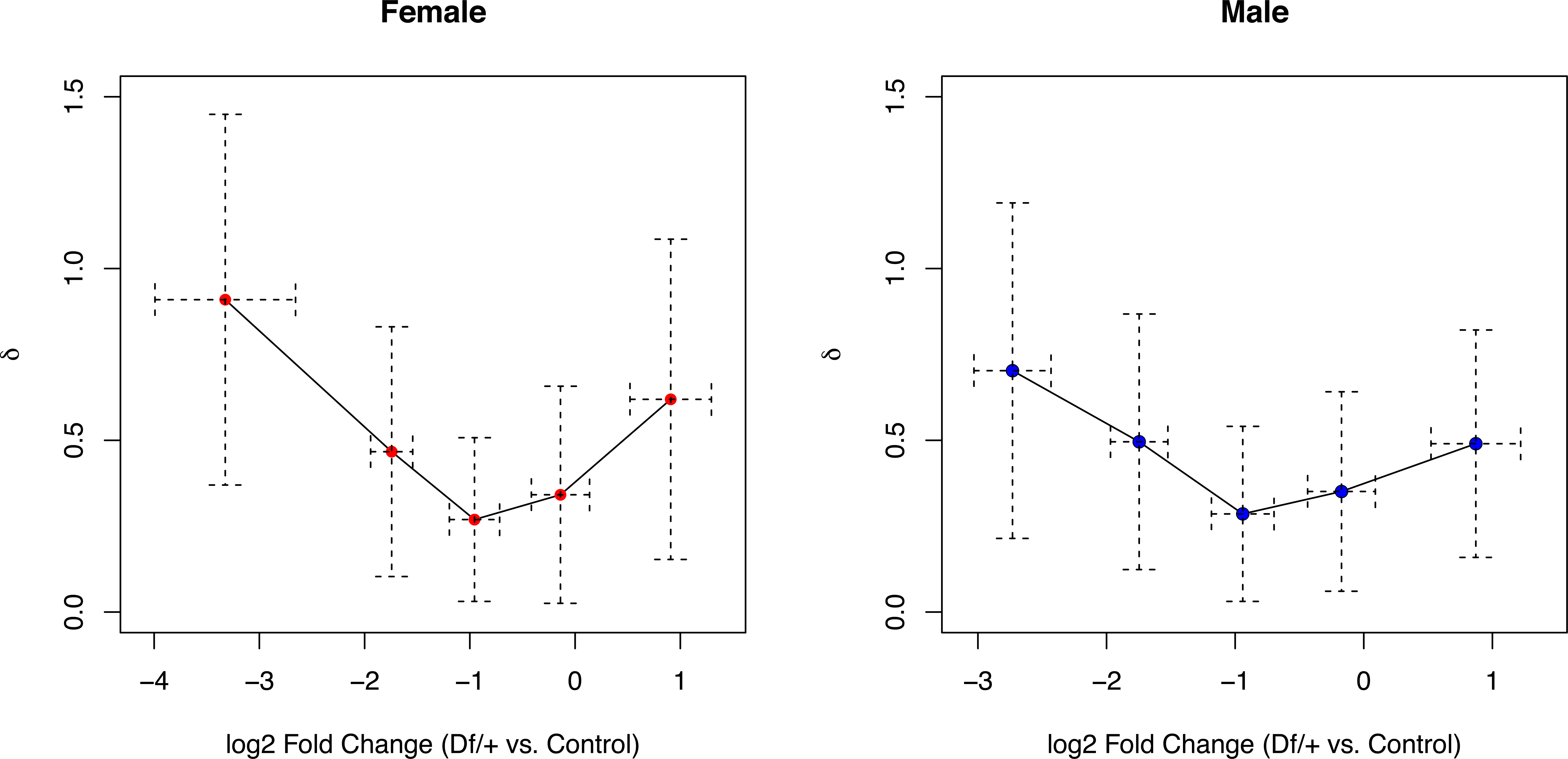
Dosage compensation vs. expression variation. Mean (dot) and standard deviation (dashed line) of δ and fold change in expression are shown for each group of one dose genes classified according to compensation levels. One copy expression baseline corresponds to log2 fold change equal to −1.

### Propagation of expression variation through gene networks

The results described above imply compensation level dependent extrinsic expression variation of one dose genes. We postulated that we should observe propagation of this extrinsic noise through the regulatory network, similar to what has been demonstrated for propagation of intrinsic noise [22,23]. That is, genes regulated by extrinsically noisy transcription factors are more likely to show extrinsic noise as well. In addition, we have previously observed that dosage compensation is, in part, mediated by interactions with other genes [20]. Therefore, we tested if the expression variation of one dose genes was propagated through the gene network. If extrinsic noise in regulators drives variation in large populations of cells, then noisy expression of regulators would propagate noise downstream. To explore this, we examined expression variation patterns through two separate paths in the gene network: one starting from one dose transcription factors (TFs) and the other from one dose non-TFs.

After projecting δ values onto a pre-existing network model [24], we examined expression variations of two dose neighbors at different network distances (Fig 4A). We found that expression variation of two dose genes gradually decreased as network distance from one dose TFs increased. In contrast, expression variations of two dose network neighbors of non-TF one dose genes were similar regardless of distance in the network model. It is also worth noting that expression variations caused by one dose TFs remain higher than expression variations unrelated to deletions in the TF, even in distant neighbors. This indicates that dose reductions in TF genes causes more genome-wide expression variation, while dose reduction of non-TF genes does not, on average, contribute significantly to global expression variance. An example of such a difference in two particular DrosDel lines is shown in Fig 4B. Males heterozygous for *Df(2L)ED1231* have one dose of two TFs (*brat* and CG17568), and show much higher global gene expression variance than males heterozygous for *Df(2L)ED279*, which uncovers no TF encoding genes. Both deletions uncover 33 genes and are of similar size (305Kb and 249Kb). These findings indicate that variability in expression of TFs is the main source of expression variation in downstream genes in the same pathway. This is consistent with the expectation that expression variability in transcription factors contribute to extrinsic noise of downstream genes. Taken together, our results show that that gene dosage related expression variability of key regulators can propagate through the network potentially contributing to the heterogeneity of phenotypic consequences that reductions in gene dosage can have on human development and physiology as revealed through haploinsufficiency-associated diseases [25,26].

**Fig 4.**
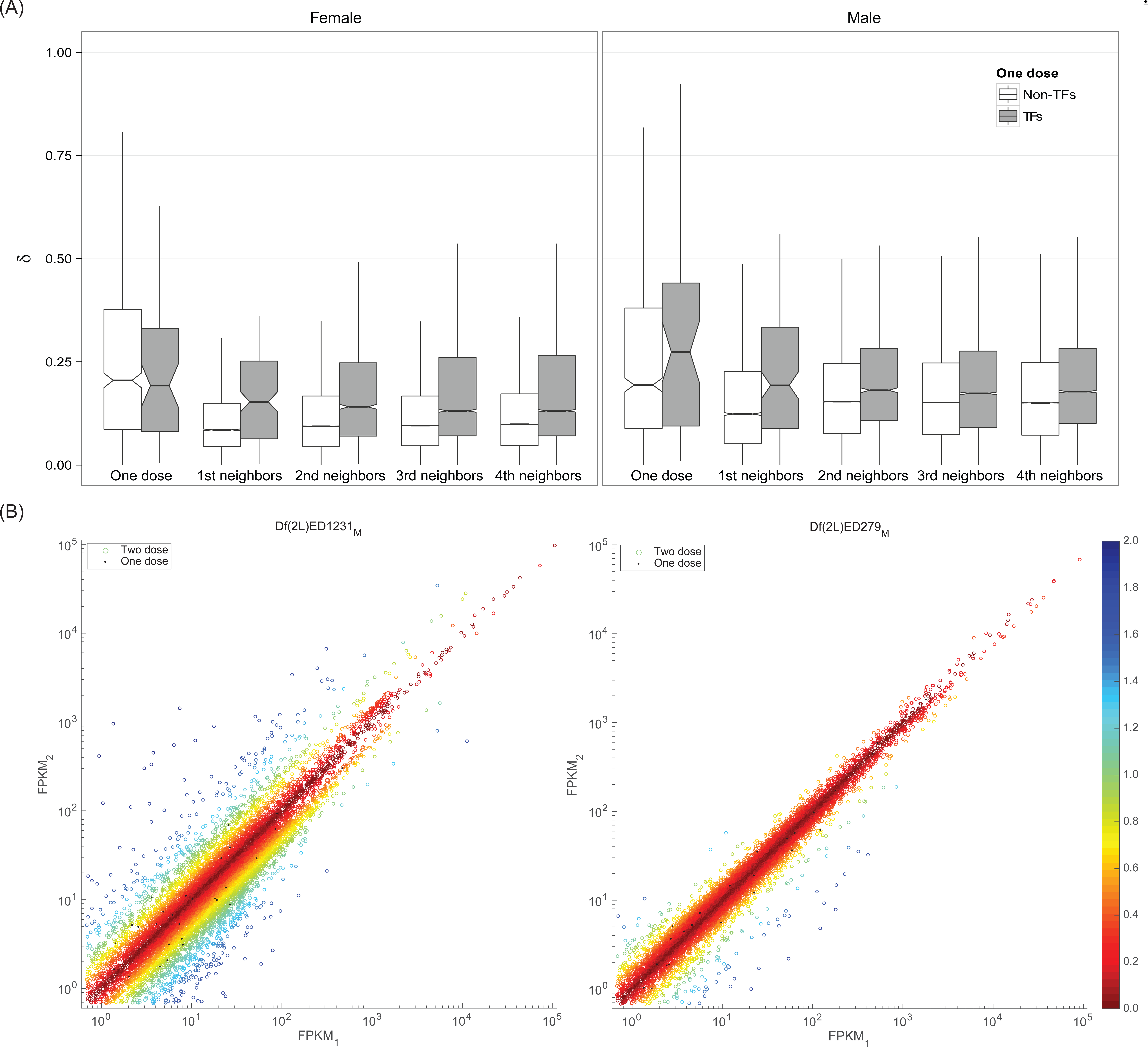
Propagation of expression variation through a network. (A) Boxplots show medians (bar), 95% confidence intervals (notch), 25 to 75 percentiles (box), and 1.5 × interquartile ranges (whisker) of the distribution of delta values for one-dose TFs (white) and non-TFs (gray), and their two dose neighbors in the functional network. Outliers are omitted. (B) Scatterplots show FPKM values for all expressed genes in two illustrative lines. The main difference between two lines is that two TFs (*brat* and CG17568) are located in *Df(2L)ED1231* deficiency (left), but all one dose genes are non-TFs in *Df(2L)ED279* (right). Both deletions uncover 33 genes and are approximately the same size (305kb and 249kb respectively). Color map represents δ values.

## Discussion

Gene expression is subject to intrinsic and extrinsic noise. Computational models indicate that a reduction in gene copy number increases intrinsic expression noise, however, the effect of one copy genes on extrinsic expression variability in diploid metazoans has not been clear. Measuring gene expression in a set of DrosDel lines carrying heterozygous deletions in a common genetic background, we found that in *Drosophila melanogaster* one dose genes show elevated levels of expression variability relative to two dose genes at the organismal level. Using simulations, we demonstrated that this increased variability cannot simply be attributed to intrinsic noise as has been previously suggested [14]. Organismal level expression variability is typically attributed to differences in genetic background, however, in our experiments the same genetic background was maintained. In the absence of genetic variations, such extrinsic signal could have originated from intrinsic single cell noise during development or cell-to-cell communication. Indeed, adult tissues are derived from a very small founder population. It is thus possible that intrinsic noise in founder cells impacts early development and, subsequently, causes extrinsic gene expression in progeny cells.

Interestingly, there is a relationship between dosage compensation and expression variability. We previously observed that *Drosophila melanogaster* displays a marked heterogeneous response to copy number reduction of autosomal genes [20,21]. Many one copy genes show some dosage compensation via increased gene expression, however, levels of compensation vary drastically for individual genes. This variation can potentially be related to the differences in the need for compensation and the ability of cell circuitry (i.e., gene regulation) to deliver corresponding compensation. In contrast, *X*-linked genes in males are precisely compensated using an MSL-complex dependent mechanism [17–19]. The low extrinsic noise of one copy *X*-linked genes in males relative to those same one copy genes in *XX* females, or *XX* sex transformed flies, suggests that the MSL mechanism ensures that compensation of *X*-liked genes is precise in both compensation level and expression consistency. In stark contrast, for autosomal gene deletions we observed the smallest expression variation in genes that are not compensated and highly compensated genes showed increased expression variation. Thus, our results suggest that when autosomal dosage compensation is present, regulatory circuitry is unable to provide a consistent compensation. This inability to provide a consistent expression response might be related to a lack of evolutionary pressure tuning the feedback mechanism to provide precise responses to relatively rare evolutionary events such as gene deletion.

Gene dosage compensation has to be considered in the context of gene networks [20,27]. Analysis of intrinsic noise demonstrated that expression noise can propagate through the gene network and certain network topologies can modulate noise levels [22,23,28–31]. In particular, it has been demonstrated that negative autoregulation has the ability to reduce gene expression noise [29,32,33] and can linearize the dose-response before saturation [34]. Another recently discovered noise-reducing mechanism includes miRNA regulation [35,36]. However, such noise regulation, if indeed acting on a given gene, is most likely tuned by evolutionary forces [37–39] to buffer expression variation within a normal cell and not a cell altered by gene copy number reduction. This might explain why gene dosage compensation is not only incomplete but also noisy.

A recent study in humans [40] modeling the link between copy number variantions and haploinsufficieny found that genes predicted to exhibit phenotypes when present in one copy were enriched for TFs and that the best predictor of haploinsufficieny of a gene is its proximity in probabilistic gene networks to haploinsufficient genes. Our results show the success of such predictor can be explained by the fact that gene dosage reduction in TF encoding genes is associated with a marked increase in gene expression variability propagating through the gene network.

In humans, copy number variations are associated with several complex disorders [41,42] including a variety of neuropsychiatric disorders such as autism [43,44]. Such disorders are phenotypically heterogeneous and are assumed to be caused by a combination of genetic and environmental factors. The identification of genetic causes in complex diseases is particularly challenging since genetic factors in affected individuals are often different. In recent years, network based approaches showed that these different causes are not unrelated but rather dysregulate the same component of the cellular system (reviewed in [45]). While these approaches help to explain why different perturbations cause the same phenotype, they do not explain why seemingly the same perturbations can show different phenotypes.

Here we show that the extrinsic expression noise in one dose TFs readily propagate across the regulatory network, affecting expression variation of connected two dose genes, and thus have the potential to affect entire cellular pathways and consequently the phenotype of an organism. Thus, the response to autosomal gene deletion introduces a perturbation that may influence gene network neighbors, not only through reduced expression level, but also through an increase in expression variation. These variations have the potential to differentially perturb gene subnetworks in different individuals. Thus, some of the phenotypic heterogeneity in diseases associated with gene deletions may be related to extrinsic variation in gene expression of one dose gene(s).

## Materials and methods

### Flies and RNA-Seq

The Drosophila deletion stocks were generated by DrosDel (http://www.drosdel.org.uk) as described in [15]. Flies were maintained at 25°C on standard yeast/cornmeal medium (Fly Facility, University of Cambridge, UK). We crossed males from DrosDel lines to *w^1118^* virgin females to generate hemizygotes without balancer chromosomes. The *w^1118^* line we used is the parental line for the DroDel project, so all flies are isogenic other than for the deleted region they carry. We prepared duplicates of 15-25 sexed *Df/+* progeny (mean 21.65, standard deviation 7.4) 5 days post eclosion for RNA extraction. Flies were homogenized in 500 μl of RNAlater solution (Life Technologies, Grand Island, NY, USA) for 90 seconds in 96 deep well plates with 3 mm tungsten carbide beads (Qiagen, West Sussex, UK) using a Qiagen Tissue Lyser II (Qiagen, West Sussex, UK). Extraction of total RNA was performed using the RNeasy 96 kit (Qiagen, Valencia, CA, USA) based on the manufacturer’s guide (Protocol for Isolation of Total RNA from animal cells using QIAvac96 vacuum manifold, Cat#19504). We made a minor modification to dilute the RNAlater solution, so instead of mixing 150 μl of RLT buffer (Qiagen Cat#79216) and 150 μl of 70% ethanol, we mixed 150 μl of RLT buffer, 120 μl of 100% ethanol, and 30 μl of lysate in RNAlater and applied this to the extraction columns. Libraries were prepared using Illumina’s (San Diego, CA, USA) TruSeq RNA sample preparation kit v2 - set A and B (Cat # RS-122-200). RNA amount was quantified using RiboGreen kits (Life Technology, Grand Island, NY, USA). 100 ng of RNA was used as input to follow the 1/2 scaled protocol from the original manufacturer’s High Sample Protocol (Illumina, Part# 15026495 Rev. C). We added 10 pg (353 samples) or 500 pg (43 samples) of external RNAs as spike-in [46] during the 8 minute RNA fragmentation step. The constructed libraries were sequenced with a HiSeq 2000 (Illumina, San Diego, CA, USA). The sequencing was performed as unstranded single-end sequencing with 76 or 101 base pair read-length.

RNA-Seq reads were aligned to the Drosophila genome assembly (Berkley Drosophila Genome Project Release 5, obtained from FlyBase. http://www.flybase.org) using TopHat 2.0.10 [47] with-g 1 and-G parameters. As a gene model for the-G parameter, we used an annotation file from FlyBase (5.57) where we removed genes from uncertain physical locations (e.g. chrU and chrUextra). From the alignment result, Fragments per Kilobase per Million mapped reads (FPKM) values were calculated using Cufflinks [48]. Sequencing reads from external spike-ins were not included in Cufflinks analysis to avoid their influence on FPKM measurements. Instead, FPKM values for the spike-ins were separately calculated based on the number of raw reads mapped to the spike-in sequences. We used-G,-b, and-u parameters in running Cufflinks. In describing gene expression fold changes, abundance of transcripts were measured as read counts using HTseq [49] with default options (-m union-t exon). To obtain gene expression fold differences, we compared one deficiency line to the all other deficiency lines but itself form the same sex. This process was performed using “voom” with the R limma software package [50]. The raw sequencing data as well as their count and FPKM values are available in Gene Expression Omnibus (GEO) with the accession number GSE61509.

### Measure of expression variability

To evaluate expression variation, we used RNA-Seq data from the two biological replicates. Thus, for each gene we had two measurements of mRNA levels represented by FPKM values and we calculated the expression variation metric defined by absolute difference of two FPKM values divided by their mean:

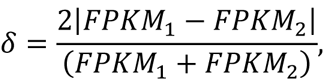

where only expressed genes in both replicates (FPKM1 and FPKM2 ≥ 0.6829118) were considered. This gene expression cutoff was made based on RNA-Seq signals from intergenic regions. We used the median value of the top 95 percentile of the intergenic signals as described in [21]. For just two measurements, δ has a linear relationship with the coefficient of variation (CV), which has been widely used as a metric of cell-to-cell expression noise:

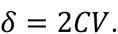

For all expressed one dose genes in the 99 DrosDel lines, Fig S3 shows the discrepancy between two replicates using our expression variation metric.

### In silico simulation

The simulation was based on a stochastic model of gene expression, in which formation and decay of single molecules occur in a random way [1]. In the model, genes function independently of each other and can switch spontaneously between repressed and active states with reaction rates k_ON_ (activation) and k_OFF_ (repression). Active genes are transcribed to mRNA with a constant rate s_A_; once a gene is activated, mRNA accumulates until the gene is deactivated. The mRNA degradation rate is δ_M_. All processes are represented by first-order kinetics reactions. Simulations were performed using STOCKS ver. 1.02 [46] software for the stochastic kinetic simulation of biochemical processes using the Gillespie algorithm [47]. All simulations started with one or two independent copies of each gene in the repressed state for one or two dose genes, respectively. The total number of mRNA copies was reported at each step of each simulation. Each independent simulation represented a time series of changes of mRNA abundance in a single cell generated from the stochastic model. Fluctuations of mRNA copies in single cells from such simulations are attributed only to intrinsic noise.

To model the expression of a single gene from a population of cells, we ran a number of independent simulations (with the same parameters) and we computed an average number of mRNA copies for each time point over these simulations. An mRNA level variation measured by δ value was computed based on two independent runs of such a computation. The computations were done for different cell population sizes. All simulations were repeated to sample the distribution of δ values for each cell population size (see Fig 2).

The simulations of one copy (Fig 2) were performed for activation and repression rates k_ON_ = k_OFF_ = 0.02/s (half-time: 35s), transcription rate s_A_ = 0.02/s and degradation rate δ_M_ = 0.008/s (half-time: 14 min); in the simulation of two copy genes, we assumed s_A_ = 0.01/s to get the same expression level as for a one copy gene. The dependence of expression variation, measured by δ, from promoter rates, transcription rates and degradation rates is presented in Fig S4.

### Compensation

To group the one dose genes with respect to their fold changes in expression relative to two dose reference (see Flies and RNA-Seq), we rounded fold changes to the nearest integer and defined five groups according to the integers:

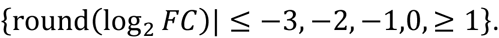

For each group, we calculated means and standard deviations of fold changes and δ values.

### Network analysis

As a network model, we used a genome-wide functional network [24] that combined various types of dataset, including genetic interactions, protein-protein interactions and microarray expression. For annotation consistency all gene IDs in the network were converted to FlyBase R5.57 [48]. To do this, we discarded genes that do not have physically defined models as well as genes that are not unequivocally identifiable in the newer annotation (e.g., gene models that have been split into multiple genes). After converting IDs we obtained a list of 613 potential TFs from the predictive regulatory model [49], 287 of which are included in the functional network. Using this TF list, for every DrosDel line, we divided one dose genes into two groups: TFs and non-TFs, then separately recorded the δ values of their two dose neighbors according to geodesic distances along the functional network.

## Authors’ contributions

DYC, HL, TMP and BO conceived and directed the project. DW performed simulations. DYC, HL, DW, TMP and BO analyzed and interpreted results. DYC, HL, DW, SR, BO and TMP wrote the manuscript. All authors have read and approved the manuscript for publication.

## Acknowledgments

We thank members of the Przytycka and Oliver labs for useful discussion. This research was supported by the Intramural Research Programs of the National Library of Medicine (NLM) and National Institute of Diabetes and Digestive and Kidney Diseases (NIDDK), National Institutes of Health (NIH), and Korean Visiting Scientist Training Award (KVSTA, HI13C1282) to HL. This research utilized the high-performance computational capabilities of the Biowulf system at the NIH, Bethesda, MD. Certain commercial equipment, instruments, or materials are identified in this document. Such identification does not imply recommendation or endorsement by NIH.

## Supporting information captions

Fig S1. Relationship between expression level and expression variability. For every one dose gene from 99 DrosDel lines in females and males, average expression level and δ are shown. Pearson’s correlation coefficient and p-values are also given.

Fig S2. Expression variation accompanying *X* chromosome deletions in females. Based on data from [21], boxplots show the distribution of δ values for different copy number classes on the X chromosome and autosomes in females heads (outliers are excluded). P-values are obtained from Wilcoxon rank sum tests.

Fig S3. Discrepancy between two replicates. All one dose genes expressed in both replicates are shown (block dots). Our expression variability metric (δ) values are represented by contour lines.

Fig S4. Dependence of expression variation on promoter transition rates, transcription rates and degradation rates. Intrinsic noise in simulated gene expression model dissolves with increasing size of cell population - dependence of gene variation from parameters of stochastic model. Boxplots represent distribution of simulated expression variation as measured by δ value for increasing size of cell population (number of simulations) per replicate in a stochastic gene model. The reference simulation (white) was performed using activation and repression rates k_ON_ = k_OFF_ = 0.02/s (half-time: 35s), transcription rate s_A_ = 0.02/s and degradation rate δ_M_ = 0.008/s (half-time: 14min). The comparison was done to show how the results depended on (A) an increase of promotor transition rates (k_ON_ = k_OFF_ increased by 10 and 100 fold), (B) an increase of transcription rate (s_A_ increased by 10 and 100 fold), and (C) a proportional increase of transcription and degradation rates (s_A_ and δ_M_ proportionally increased by 10 and 100 fold). Simulations were performed using stochastic kinetic simulation of biochemical processes with the Gillespie algorithm as described in Materials and methods.

